# A single chromosome assembly of *Bacteroides fragilis* strain BE1 from Illumina and MinION nanopore sequencing data

**DOI:** 10.1101/024323

**Authors:** Judith Risse, Marian Thomson, Garry Blakely, Georgios Koutsovoulos, Mark Blaxter, Mick Watson

## Abstract

**Background:** Second and third generation sequencing technologies have revolutionised bacterial genomics. Short-read Illumina reads result in cheap but fragmented assemblies, whereas longer reads are more expensive but result in more complete genomes. The Oxford Nanopore MinION device is a revolutionary mobile sequencer that can produce thousands of long, single molecule reads.

**Results:** We sequenced *Bacteroides fragilis* strain BE1 using both the Illumina MiSeq and Oxford Nanopore MinION platforms. We were able to assemble a single chromosome of 5.18 Mb, with no gaps, using publicly available software and commodity computing hardware. We identified gene rearrangements and the state of invertible promoters in the strain.

**Conclusions:** The single chromosome assembly of *Bacteroides fragilis* strain BE1 was achieved using only modest amounts of data, publicly available software and commodity computing hardware. This combination of technologies offers the possibility of ultra-cheap, high quality, finished bacterial genomes.

## Background

*Bacteroides fragilis* is a gram-negative, obligate anaerobic bacterium that is commensal in the human colon; however it is also an opportunistic pathogen and is a major cause soft tissue infections. The *B. fragilis* lipopolysaccharide (LPS) triggers an inflammatory immune response *via* the Toll-like receptor 2 (TLR2) [1] pathway. Significant intra-strain antigenic variation has been observed, suggested to be an adaptation to its survival in the human host [2]. The first *B. fragilis* genomes (NCTC 9343 and YCH46) were sequenced in 2004-2005 [3, 4]. These projects identified dynamic rearrangement in *B. fragilis*, including several invertible promoters associated with LPS biosynthesis gene clusters and four “shufflons” that had the potential to alter gene expression of specific genes. Together, these inversions regulate cell surface adaptation and bacterial phage resistances and are the source of the observed antigenic variation [4].

Second-and third-generation sequencing instruments are revolutionising biology and medicine [5]. Cheap “benchtop” instruments enable access to huge sequencing power even for smaller laboratories [6], and instruments such as Illumina’s MiSeq are capable of sequencing millions of 600 base fragments simultaneously. Illumina’s higher-throughput sequencers produce up to 1.8 terabases of sequence per run [7]. The throughput of these machines has enabled scientists to sequence thousands of bacterial genomes at low cost [8]. However, due to the relatively short reads and insert lengths, genome assemblies from Illumina data tend to be fragmented, because the read and insert lengths are shorter than repeat regions within the genome. The problem of fragmented short-read assemblies has led many researchers to use Pacific Biosciences (PacBio) sequence data to assemble bacterial genomes, often (but not always) resulting in chromosome-level assemblies [9]. PacBio sequencing uses a modified DNA polymerase and produces long (∼17 kb) single-molecule reads with a high individual error rate, which can be corrected to high accuracy [9, 10]. Whilst PacBio assemblies are of higher quality, they come at approximately 3-4 times the cost [8].

The Oxford Nanopore MinION is new mobile sequencing machine. The size of an office stapler, the device is powered by the USB port of a laptop computer. The MinION measures changes in the electronic current as single molecules of DNA are passed through a biological nanopore. By using a hairpin adapter, each molecule is read twice and the resulting 2D reads are long (usually 5-6 kb, but there is no theoretical limit) [11–13]. The first nanopore-only bacterial genome assembly has been published [14]. However the assembly process was complex and the resulting assembly has a high error rate (1,202 mismatches and 17,241 indels). Others have reported using MinION data to successfully arrange Illumina contigs into a single scaffold [15].

At time of publication [16], there are 103 registered *Bacteroides fragilis* genome projects; however, only four are listed as complete, including those mentioned above, strain 638R and strain BOB25 [17]. As part of our Junior Honours “Genomes and Genomics 3” undergraduate course, we run a practical for ∼100 students in bacterial genome sequencing, annotation and analysis, and the class of 2013 sequenced eight previously unanalysed *B. fragilis* strains using Illumina MiSeq. The assemblies generated were, as expected, fragmented, and it was not possible to definitively map polysaccharide biosynthesis clusters.

Here we present a fully contiguous, single-chromosome assembly (with no gaps) of *Bacteroides fragilis* strain BE1, a previously unsequenced strain originally isolated from the wound infection of a patient at the Academic Hospital of the Vrije Universiteit [18]. The assembly was produced using open-source tools and a combination of Illumina MiSeq and MinION nanopore data. Crucially, the finished genome was achieved using only moderate amounts of data and assembled on commodity computing hardware, suggesting that high-quality, finished bacterial genomes can be achieved at very low cost with only a small amount of bioinformatics infrastructure.

## Results

### Sequence data characteristics

The MiSeq data, generated by the Genomes and Genomics 3 class, consisted of 898,420 2x250bp reads. After adapter removal and trimming for low quality the reads had a mean length of 248bp. The MinION run produced 7300 2D MinION reads with a mean length of 6618 and maximum length of 29,630 (Figure 1).

**Figure 1.**
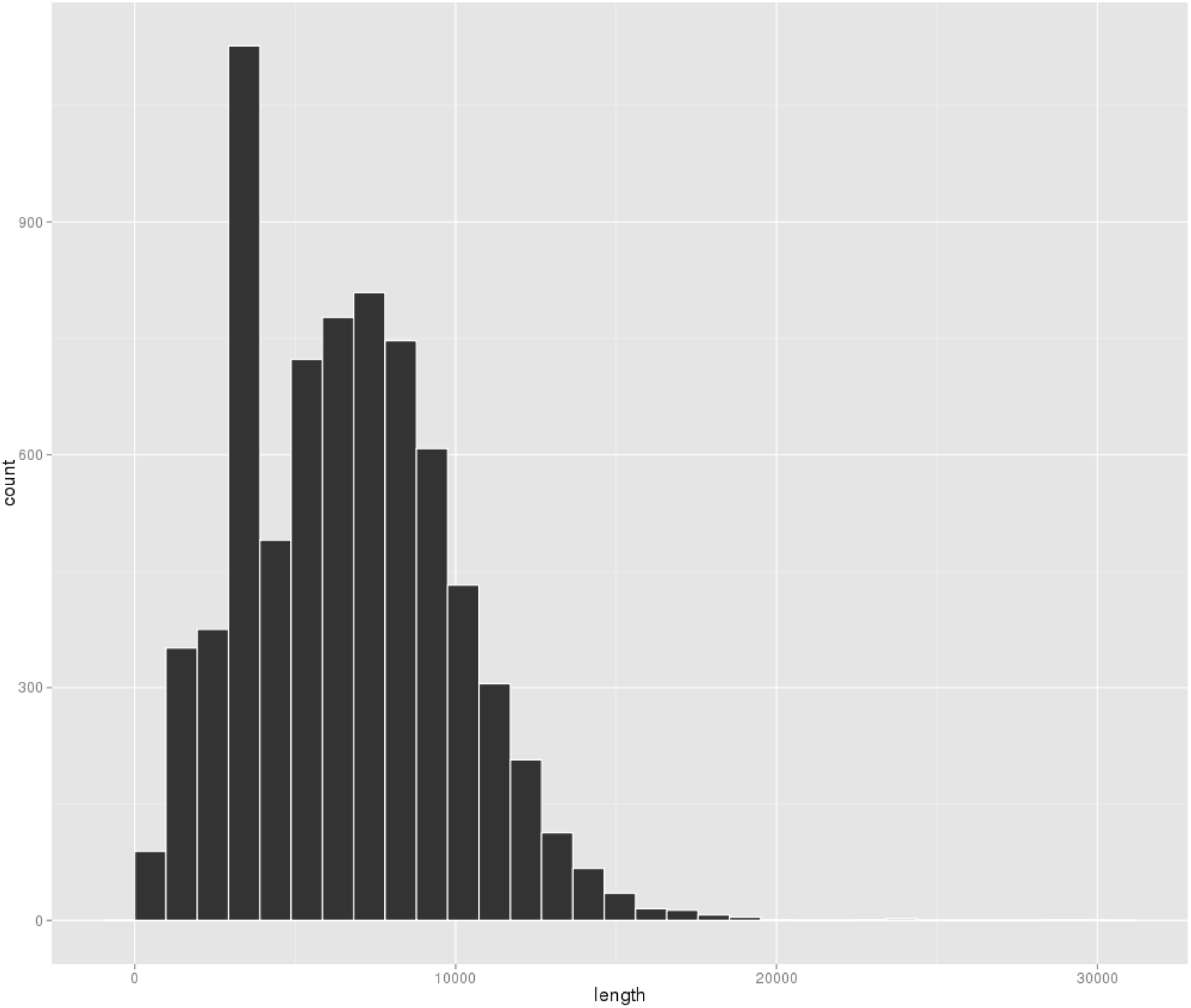
MinION read-length histogram. A histogram of 2D read lengths from 7300 MinION reads. The peak at 3-4kb represents the lambda spike-in

### Assembly and genome characteristics

The complete genome of *Bacteroides fragilis* strain BE1 has a length of 5,188,967 base-pairs and a GC content of 43.1%, consistent with other strains. Genome annotation identified 4217 coding sequences (CDS), 18 rRNA genes and 74 tRNA genes.

Post-assembly assessment showed that 99.16% of the MiSeq reads mapped to the assembly and 98.87% were marked as properly paired. For the MinION reads, 6640 (88.2%) mapped to the *B. fragilis* BE1 assembly while the remaining 830 (11.8%) mapped to the phage lambda genome, used as a spike-in during library preparation. MinION percentage identity to the assembly (calculated as *100 * matches/(matches + deletions + insertions + mismatches*)) is an average of 85% (standard deviation: 2.64) (Figure 2). The 2D alignment lengths were all approximately equal to the read length, albeit with a slight tendency for the alignment length to be greater than 2D sequence length (Figure 3). The high mapping rate of properly paired Illumina reads, in combination with the high number of mapped MinION reads is indicative of a high quality genome assembly. The entire assembly is covered by at least one full length 2D MinION read. The average coverage from MinION data is 8.7X.

**Figure 2.**
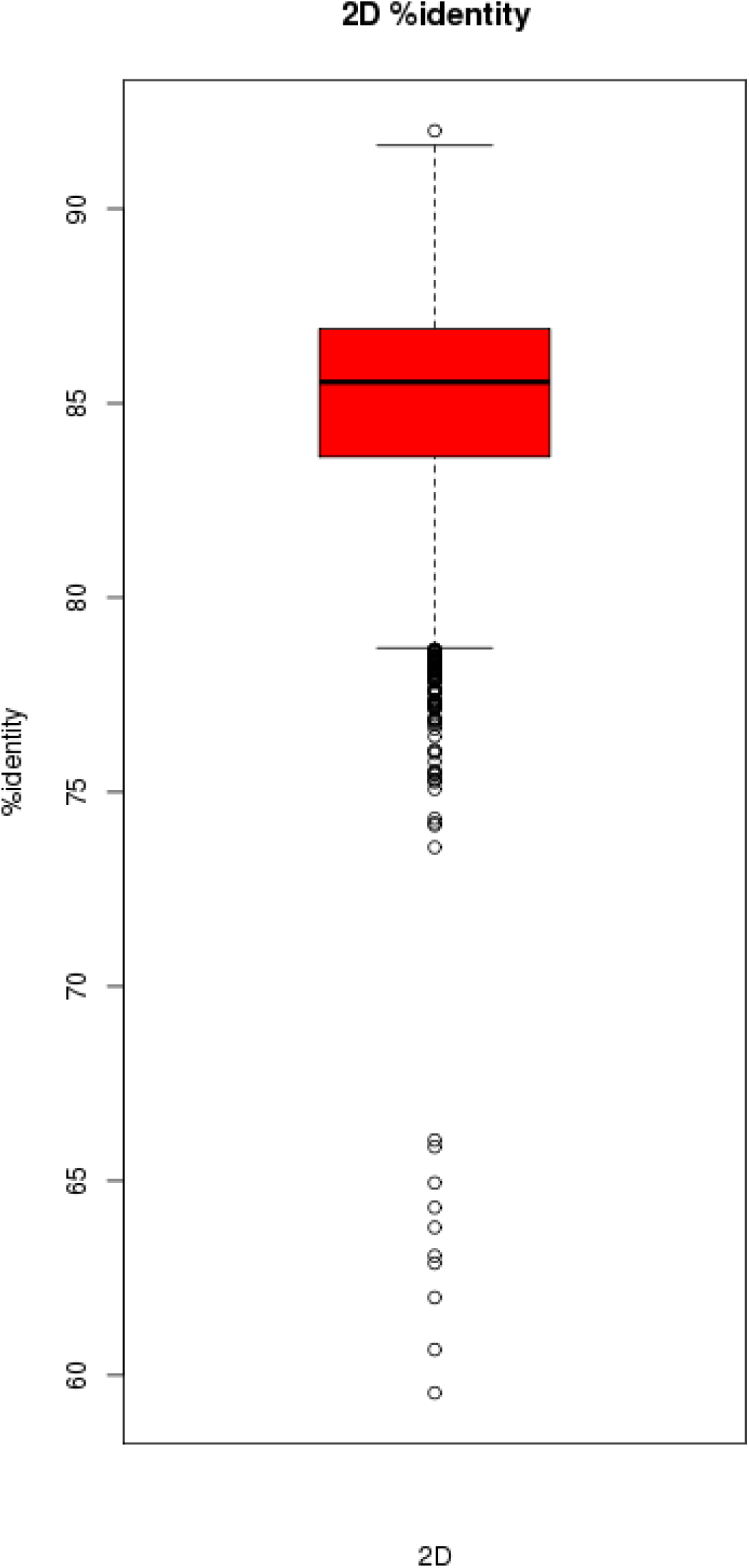
MinION percentage identity. A boxplot of percentage identity to our assembly for 7300 MinION reads

**Figure 3.**
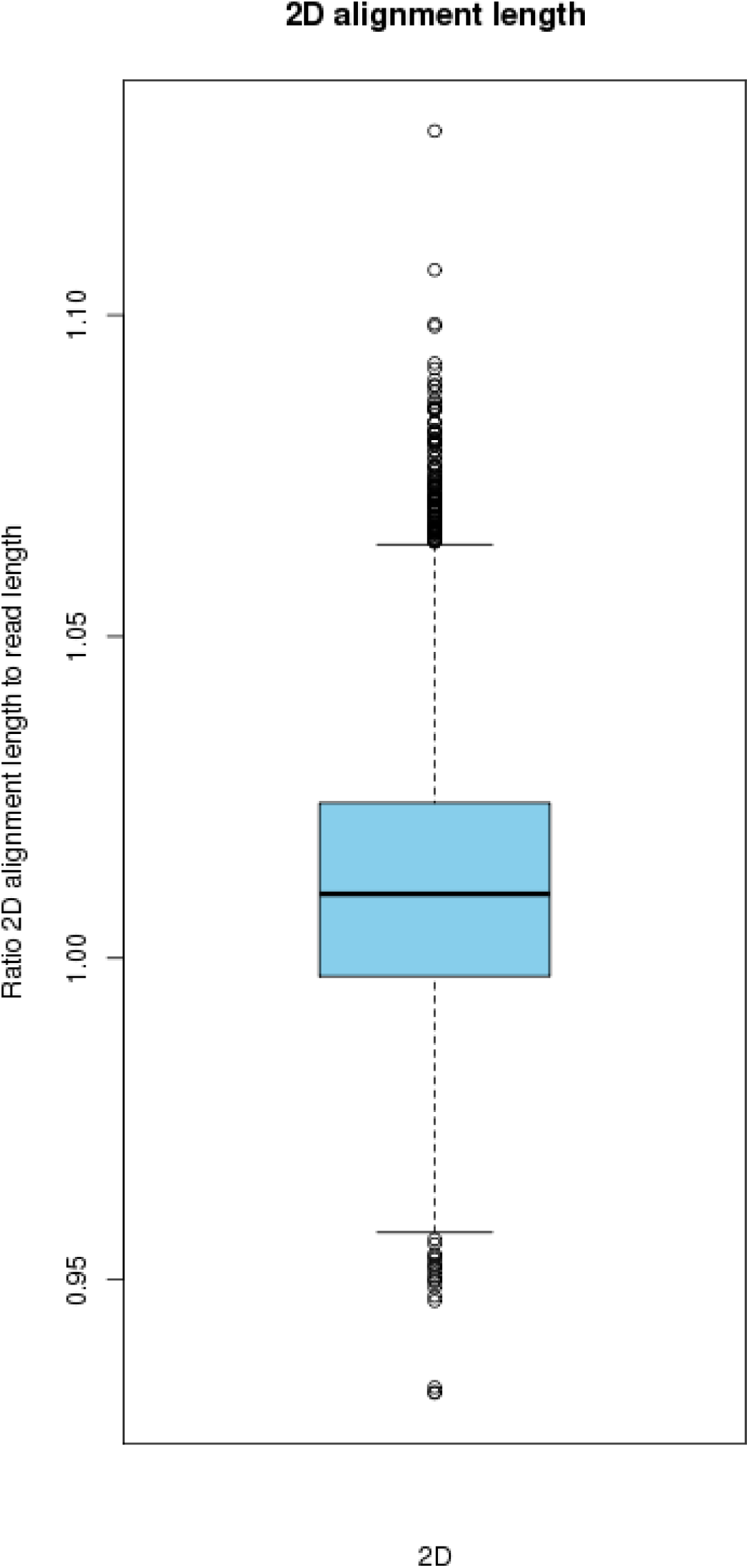
MinION alignment length. A boxplot showing the ratio of alignment length to read length for 7300 MinION reads

In addition to the internal, read-mapping consistency, regions of difference (RODs) between our assembly and NC_003228 were manually inspected to assess assembly integrity; specifically we checked that presence/absence of the RODs were supported by the read data. We identified several regions where our assembly evidently has a small inversion compared to the reference genomes, and these correlated with invertible promoters

### Data availability

Raw Illumina and MinION reads and the annotated assembly are available in the European Nucleotide Archive under project accession PRJEB10044.

## Discussion and Conclusions

*Bacteroides fragilis* is a commensal bacterium of the human colon; however, it is also an opportunistic pathogen, being one of the major causes of soft-tissue infections in humans. Significant intra-strain antigenic variation has been observed in *B fragilis,* caused by promoter DNA inversions that regulate gene expression of cell surface antigens.

Second and third-generation sequencing technologies now enable the rapid and accurate sequencing of thousands of bacterial genomes. Indeed, these technologies are easily accessible to even undergraduate practical classes, giving students direct experience of genomics in practice. However, assemblies created using only Illumina “short reads” are often fragmented. Therefore, long read sequence data (e.g. PacBio) is now regularly used to finish and complete bacterial genomes. The Oxford Nanopore MinION is the world’s first mobile DNA sequencer, capable of producing long, single-molecule reads, and the aim of this study was to discover whether MinION reads could be used to finish and complete a bacterial genome.

Here we describe the complete, finished genome of *Bacteroides fragilis* strain BE1 using a combination of Illumina and Oxford Nanopore MinION data. To our knowledge, this is the first new bacterial genome finished using a combination of Illumina and MinION nanopore data. The high quality Illumina data in combination with long MinION reads has resulted in a fully circularised, contiguous, and high quality assembly.

Crucially, the assembly was created using free, open-source bioinformatics tools, on commodity computing hardware (16 cores; 64Gb RAM) using only a moderate amount of data. The data volumes used are modest: 898,420 MiSeq reads is approximately 8% of a MiSeq V2 run, and 7300 MinION reads is approximately 20% of a MinION run. Assuming a 2x250 MiSeq run costs £1400 and a MinION run costs £800 (approximate full economic costs from Edinburgh Genomics [20]), the sequence generation costs would be approximately £276 per genome. Even when adding library preparation costs, it is easy to imagine that a fully circularised, complete bacterial genome could cost less than £500. Given the low capital expenditure associated with MiSeq and/or MinION sequencers, we predict that Illumina+MinION bacterial genome sequencing will become the norm in the short-to medium-term future.

## Methods

### Strain growth and DNA extraction

*B. fragilis* was grown in a Don Whitley Scientific (UK) MiniMacs anaerobic work station at 37°C with an anaerobic gas mix (10% hydrogen, 10% carbon dioxide and 80% nitrogen), in brain heart infusion broth (BHI) (Difco, USA) supplemented with 5% cysteine, 10% sodium bicarbonate, 50 μg/ml haemin and 0.5 μg/ml menadione. DNA was extracted from stationary phase cultures of *B. fragilis* using the Promega Wizard Genomic DNA Purification Kit (as per manufacturer’s instructions), and secondarily cleaned of residual RNA using Riboshredder (Epicenter, USA) and Zymoclean (Zymoresearch, USA) columns. DNA was quantitated using Qubit (Life Technologies, UK).

### Illumina library construction and sequencing

1 ng of input DNA was simultaneously fragmented and tagged with specific Illumina adapter sequences by the Nextera XT transposome complex, as described in the Nextera XT DNA Library Preparation protocol (illumina). Following a neutralisation step, the sample, was amplified by limited cycles of PCR, which also added sequencing primer sequences to tagmented DNA fragments. The library was then prepared for cluster generation, and sequenced on a Miseq (Illumina) 250 base paired-end run.

### MinION library construction and sequencing

Library preparation was carried out using the Nanopore Genomic Sequencing Kit (SQK-MAP005) and following Version *MN005_1124_revC_02Mar2015* of the Oxford Nanopore protocol. After extraction, the DNA was purified by Agencourt AMPure XP beads (Beckman Coulter Inc) at 1.8:1 bead to DNA ratio, and quantified by Qubit High Sensitivity assay (Life Technologies). 2 μg of DNA was sheared in a total volume of 80 μl Tris Cl pH 8.5 by G-tube (Covaris) centrifugation at 5200 rpm (Heraeus Pico21 Thermo Scientific) for 60 s, followed by a repeat 5200 rpm 60 s spin after inversion of the G-tube. The resultant fragment size distribution was determined by DNA 12000 Bioanalyzer assay (Agilent Technologies Inc), and the recovered DNA was re-quantified by Qubit. To minimise the effect of potential DNA damage on sequencing library performance, 1 μg of the sheared DNA was repaired using the PreCR Repair Mix (New England BioLabs) prior to commencement of library preparation.

To prepare the DNA for MinION sequencing, the DNA was first end-repaired and then dA tailed using NEBNext End-Repair, and NEBNext dA-Tailing Modules (New England BioLabs) according to manufacturer’s instructions. Each reaction was cleaned, and smaller fragments excluded using Agencourt AMPure XP beads at 0.5:1 bead to DNA ratio. Specific adapters (SQK-MAP005; Oxford Nanopore) were then ligated to the dA-tailed DNA using Blunt/TA Ligase Master Mix (New England BioLabs). These adapters comprise: a leader adapter responsible for movement of DNA through the pores, and a “hairpin” adapter which links the 2 strands of the DNA molecule and permits sequencing of both DNA strands (2D reads). One of adapters is also His-tagged, enabling selection of adapter ligated fragments by His-Tag Isolation and Pulldown Magnetic Dynabeads (Life Technologies) following version *N005_1124_revC_02Mar2015* protocol guidelines.

The library eluted from the beads and was quantified by Qubit prior to sequencing on a MinION device (original “Mark 0”). An R7.3 FLO-MAP003 flowcell was attached to the MinION, connected to a laptop via a USB port. Platform QC was first carried out to determine the number of viable pores available for the sequencing run. The flow cell was primed with sequencing buffer then 220ng of freshly prepared library diluted in sequencing buffer was added to the flowcell via the sample port. A 48-hour gDNA sequencing run was initiated using the MinION™ control software, MinKNOW version 0.49.3.7 and the run was topped up with diluted library at 12 hour intervals.

### Assembly and annotation

Illumina reads were trimmed using Trimmomatic [21]. Sequencing adapters were removed, as were bases less than Q20. Any reads less than 126bp in length after trimming were discarded. MinION reads were extracted using poRe [22]. Input data for the assembly were therefore 898,420 250 base paired-end Illumina MiSeq reads and 7300 2D MinION reads with a mean length of 6618 and maximum length of 29,630. These were used as input to SPAdes [23] version 3.5.0 with the 

~~~
--nanopore
~~~

 option.

After removal of short and/or low-coverage contigs, the SPAdes hybrid assembly consisted of 5 contigs of length 3980468, 827237, 362398, 13363, 5146 nucleotides respectively. The smallest contig had a reported coverage 6-times that of the other 4 and contained an rRNA operon, suggesting that there are 6 copies of that operon within the genome.

These 5 contigs were used as input to SSPACE-LongRead [24] using the MinION reads to scaffold, resulting in 3 scaffolds of length 48125973, 362398, 13363 nucleotides respectively. This scaffolding step placed the rRNA operon successfully into 6 locations in the larger scaffolds. The three scaffolds were used as input to a second round of SSPACE-LongRead which produced a single scaffold of length 5188967 and containing 3 gaps. These gaps were successfully filled using GapFiller [25] and the paired-end Illumina data.

The chromosome start was defined by comparison to sequence NC_003228 (*Bacteroides fragilis* NCTC 9343) and annotated using Prokka v 1.11 [26].

### Read mapping

Illumina and MinION reads were mapped back to the final assembly using bwa mem v0.7.12 [19], and the resulting alignments converted to BAM, indexed and sorted using Samtools [27]. MinION reads were mapped using the option “-x ont2d” in bwa mem. MinION reads were also mapped using last [28] and parameters -q 1 -a 1 -b 1. Mapping statistics were calculated using count-errors.py [29], modified slightly to work with our read IDs.

## Authors’ contributions

JR carried out bioinformatics analysis and helped draft the paper.

MT carried out MinION sequencing and helped draft the paper.

GB conceived the study, grew the *B. fragilis* cells, organised the GG3 class, extracted DNA and helped draft the paper.

GK carried out bioinformatics analysis and helped draft the paper.

MB conceived the study, organised the GG3 class, and helped draft the paper.

MW carried out bioinformatics analysis and helped draft the paper.

## Acknowledgements

The authors would like to thank Oxford Nanopore for granting Edinburgh Genomics access to the MinION Access Programme (MAP).

The DNA preparation and library construction for MiSeq sequencing of BE1 was carried out by members of the Junior Honours “Genomes and Genomics 3” undergraduate class in the School of Biological Sciences, to whom we offer fullsome thanks. The work was enabled by funding from the Biotechnology and Biological Sciences Research Council including Institute Strategic Programme and National Capability grants (BBSRC; BBS/E/D/20310000, BB/J004243/1, BB/M020037/1). Edinburgh Genomics is partly supported through core grants from the National Environmental Research Council (NERC R8/H10/56), Medical Research Council (MRC MR/K001744/1) and The Biotechnology and Biological Sciences Research Council (BBSRC BB/J004243/1).

## References

1. Erridge C: Lipopolysaccharides of Bacteroides fragilis, Chlamydia trachomatis and Pseudomonas aeruginosa signal via Toll-like receptor 2. J Med Microbiol 2004, 53:735–740.

2. Patrick S, Gilpin D, Stevenson L: Detection of Intrastrain Antigenic Variation of Bacteroides fragilis Surface Polysaccharides by Monoclonal Antibody Labelling. Infect Immun 1999, 67:4346–4351.

3. Cerdeño-Tárraga AM, Patrick S, Crossman LC, Blakely G, Abratt V, Lennard N, Poxton I, Duerden B, Harris B, Quail MA, Barron A, Clark L, Corton C, Doggett J, Holden MTG, Larke N, Line A, Lord A, Norbertczak H, Ormond D, Price C, Rabbinowitsch E, Woodward J, Barrell B, Parkhill J: Extensive DNA inversions in the B. fragilis genome control variable gene expression. Science 2005, 307:1463–5.

4. Kuwahara T, Yamashita A, Hirakawa H, Nakayama H, Toh H, Okada N, Kuhara S, Hattori M, Hayashi T, Ohnishi Y: Genomic analysis of Bacteroides fragilis reveals extensive DNA inversions regulating cell surface adaptation. Proc Natl Acad Sci U S A 2004, 101:14919–24.

5. Watson M: Illuminating the future of DNA sequencing. Genome Biol 2014, 15:108.

6. Loman NJ, Misra R V, Dallman TJ, Constantinidou C, Gharbia SE, Wain J, Pallen MJ: Performance comparison of benchtop high-throughput sequencing platforms. Nat Biotechnol 2012, 30:434–9.

7. Welcome to the $1,000 genome: on Illumina and next-gen sequencing - Biome [http://biome.biomedcentral.com/welcome-to-the-1000-genome/]

8. Koren S, Harhay GP, Smith TPL, Bono JL, Harhay DM, Mcvey SD, Radune D, Bergman NH, Phillippy AM: Reducing assembly complexity of microbial genomes with single-molecule sequencing. Genome Biol 2013, 14:R101.

9. Chin C-S, Alexander DH, Marks P, Klammer AA, Drake J, Heiner C, Clum A, Copeland A, Huddleston J, Eichler EE, Turner SW, Korlach J: Nonhybrid, finished microbial genome assemblies from long-read SMRT sequencing data. Nat Methods 2013, 10:563–9.

10. Koren S, Schatz MC, Walenz BP, Martin J, Howard JT, Ganapathy G, Wang Z, Rasko DA, McCombie WR, Jarvis ED, Adam M Phillippy: Hybrid error correction and de novo assembly of single-molecule sequencing reads. Nat Biotechnol 2012, 30:693–700.

11. Jain M, Fiddes IT, Miga KH, Olsen HE, Paten B, Akeson M: Improved data analysis for the MinION nanopore sequencer. Nat Methods 2015, 12:351–356.

12. Loman NJ, Watson M: Successful test launch for nanopore sequencing. Nat Methods 2015, 12:303–4.

13. Urban JM, Bliss J, Lawrence CE, Gerbi SA: Sequencing Ultra-Long DNA Molecules with the Oxford Nanopore MinION. Cold Spring Harbor Labs Journals; 2015.

14. Loman NJ, Quick J, Simpson JT: A complete bacterial genome assembled de novo using only nanopore sequencing data. Nat Methods 2015, advance on.

15. Karlsson E, Lärkeryd A, Sjödin A, Forsman M, Stenberg P: Scaffolding of a bacterial genome using MinION nanopore sequencing. Sci Rep 2015, 5:11996.

16. Organism browser - Assembly - NCBI [http://www.ncbi.nlm.nih.gov/assembly/organism/817/all/]

17. Nikitina AS, Kharlampieva DD, Babenko V V, Shirokov DA, Vakhitova MT, Manolov AI, Shkoporov AN, Taraskina AE, Manuvera VA, Lazarev VN, Kostryukova ES: Complete Genome Sequence of an Enterotoxigenic Bacteroides fragilis Clinical Isolate. Genome Announc 2015, 3:e00450–15.

18. Verweij WR, Namavar F, Schouten WF, MacLaren DM: Early events after intra-abdominal infection with Bacteroides fragilis and Escherichia coli. J Med Microbiol 1991, 35:18–22.

19. Li H: Aligning sequence reads, clone sequences and assembly contigs with BWA-MEM. 2013:3.

20. Edinburgh Genomics [https://genomics.ed.ac.uk/]

21. Bolger AM, Lohse M, Usadel B: Trimmomatic: a flexible trimmer for Illumina sequence data. Bioinformatics 2014, 30:2114–2120.

22. Watson M, Thomson M, Risse J, Talbot R, Santoyo-Lopez J, Gharbi K, Blaxter M: poRe: an R package for the visualization and analysis of nanopore sequencing data. Bioinformatics 2015, 31:114–5.

23. Bankevich A, Nurk S, Antipov D, Gurevich AA, Dvorkin M, Kulikov AS, Lesin VM, Nikolenko SI, Pham S, Prjibelski AD, Pyshkin A V, Sirotkin A V, Vyahhi N, Tesler G, Alekseyev MA, Pevzner PA: SPAdes: a new genome assembly algorithm and its applications to single-cell sequencing. J Comput Biol 2012, 19:455–77.

24. Boetzer M, Pirovano W: SSPACE-LongRead: scaffolding bacterial draft genomes using long read sequence information. BMC Bioinformatics 2014, 15:211.

25. Boetzer M, Pirovano W: Toward almost closed genomes with GapFiller. Genome Biol 2012, 13:R56.

26. Seemann T: Prokka: rapid prokaryotic genome annotation. Bioinformatics 2014, 30:2068–9.

27. Li H, Handsaker B, Wysoker A, Fennell T, Ruan J, Homer N, Marth G, Abecasis G, Durbin R: The Sequence Alignment/Map format and SAMtools. Bioinformatics 2009, 25:2078–9.

28. Frith MC, Hamada M, Horton P: Parameters for accurate genome alignment. BMC Bioinformatics 2010, 11:80.

29. Github: nanopore-scripts [https://github.com/arq5x/nanopore-scripts]

